# Distinct expression profile reveals glia involvement in the trigeminal system attributing to post-traumatic headache

**DOI:** 10.1101/2024.09.30.615817

**Authors:** Gurueswar Nagarajan, Yumin Zhang

**Affiliations:** Henry M. Jackson Foundation for the Advancement of Military Medicine, 6720A Rockledge Dr, Bethesda, Maryland 20817, USA; Department of Anatomy, Physiology and Genetics, Uniformed Services University of the Health Sciences, 4301 Jones Bridge Road, Bethesda, Maryland 20814, USA

**Author notes:** Corresponding author: Yumin Zhang, MD, PhD, Dept. of Anatomy, Physiology and Genetics, Uniformed Services University of the Health Sciences, 4301 Jones Bridge Road, Bethesda, MD 20814, USA.

## Abstract

Post-traumatic headache (PTH) is a common comorbid symptom affecting at least one-third of patients with mild traumatic brain injury (mTBI). While neuroinflammation is known to contribute to the development of PTH, the cellular mechanisms in the trigeminal system crucial for understanding the pathogenesis of PTH remain unclear. Male mice underwent a non-invasive repetitive mTBI protocol (4 impacts, 24-hour intervals) targeting either the bregma or pre-bregma region of the head. Periorbital allodynia and spontaneous pain behavior were assessed using von Frey tests and grimace scores, respectively. Quantitative PCR was used to evaluate the extent of mTBI-induced pathology in several brain regions. RNA sequencing was performed to obtain transcriptomic profile of the trigeminal ganglion (TG), trigeminal nucleus caudalis (Sp5C) and periaqueductal gray (PAG) at 7 days post-TBI. Subsequently, quantitative PCR, in situ hybridization and immunohistochemistry were used to examine mRNA and protein expression of glia specific markers and pain associated molecules. The repetitive impacts at the bregma, but not pre-bregma site led to periorbital hypersensitivity, which was correlated with enhanced inflammatory gene expression in multiple brain regions. RNA sequencing identified distinct transcriptomic profiles in the peripheral TG, central Sp5C and PAG following mTBI. Using gene set enrichment analysis, positive enrichment of non-neuronal cells in the TG and neuroinflammation in the Sp5C were identified to be essential in the pathogenesis of PTH. In situ assays also revealed that gliosis of satellite glial cells in the TG and astrocytes in the Sp5C were prominent days after injury. Furthermore, immunohistochemical study revealed a close interaction between activated microglia and reactive astrocytes correlating with increased calretinin interneurons in the Sp5C. Transcriptomics analysis indicated that non-neuronal cells in peripheral TG and successive in situ assays revealed that glia in the central Sp5C are crucial in modulating headache-like symptoms. Thus, selective targeting of glia cells can be a therapeutic strategy for PTH attributed to repetitive mTBI.

## Background

Mild traumatic brain injury (mTBI) causes a myriad of pathologies and symptoms. Post-traumatic headache (PTH) is one of the underreported comorbid symptoms of mTBI [12] and is observed in over 30% of clinical diagnoses [3–11]. Repetitive head injuries are associated with an increased incidence of chronic pain, including headaches [6] and clinical data shows that the incidence of PTH varies tremendously involving different mechanisms [12]. Over the last two decades, the heterogeneous nature of PTH symptoms has led to increased understanding of different headache classifications [12]. In parallel to these classifications, there are multiple PTH phenotypes and pathology associated with the trigeminal system is thought to underlie one of the phenotypes observed in the clinical population. Several theories regarding the central sensitization that triggers headaches, including migraine, have been proposed [13–15]. However, due to profound trigeminal innervation [16], the exact cellular and molecular mechanisms through which trigeminal regions are primed after mTBI for central pain sensitization of headache remain unclear.

Sensory pseudo-unipolar neurons in the trigeminal ganglion (TG) have innervation throughout the head including the facial region, meninges, scalp, cranial bones and neurovasculature [16–19]. These sensory neurons have a tight relationship with satellite glial cells (SGC) [20,21]. Thus, neurons in the TG are prone to primary insults during head injury via the afferent fibers innervating the head and the injury may affect their first-order projections to the trigeminal nucleus caudalis (Sp5C) as well as other regions of the brainstem. However, these brain regions are subject to injury due to rotational movement [22,23] which can impact their projections to the other areas of the brain, including periaqueductal gray (PAG), thalamus, parabrachial nucleus, and anterior cingulate cortex. Thus, head injury-induced pathologies in the TG and Sp5C could cause central sensitization in chronic pain conditions [24–28].

Preclinical models of neurotrauma have been reported for the past several decades [29–31]. Due to the nature of the injury model and heterogenous symptoms associated with trauma [32–34], the underlying cellular and molecular mechanisms of trigeminal pain following head injury remain poorly understood. Recent evidence shows that invasive procedures have a major effect on sensory neurons of the TG [35,36], leading to the development of trigeminal pain [37]. Therefore, non-invasive procedures are necessary to study the trigeminal pain pathway associated with mTBI. Here we used repetitive head injury induced by the closed-head impact model of engineered rotational acceleration (CHIMERA) [38] and showed that development of headache-like symptoms depends on the localization of injury/trauma to the head. Using deep sequencing, in situ hybridization and immunohistochemistry, we aimed to understand the cellular and molecular mechanisms, particularly the inflammatory response, underlying the pathogenesis of PTH attributed to repetitive mTBI.

## Methods and Materials

### Induction of repetitive mTBI animal model

Two months old male C57BL/6-J mice were obtained from Jackson Laboratory and were habituated for a week before experimental procedures. Once habituated, mice were randomized into different groups. Baseline measurements for von Frey and grimace were obtained prior to repeated mTBI. On the day of the mTBI procedure, mice were acclimated in the procedure room for 30 min. Each mouse was anesthetized under 3% isoflurane in an induction box, and was then placed in supine position under 1.5–2% isoflurane on the CHIMERA (Closed-Head Impact Model of Engineered Rotational Acceleration) stage as described [38]. After a stabilized breathing response and within 2–3 min of anesthesia each mouse received a hit from the piston with an energy level of 0.7 J. Since the position of mTBI determines the severity of comatose and brainstem pathologies [39], mice received TBI impacts in either one of the midline positions (pre-bregma or bregma regions, *n* = 6/group) shown in Fig. 1. The distance between the two hit positions differed by ∼ 0.5 cm. Immediately following mTBI, each mouse was placed in the supine position on fresh bedding material and their righting reflexes (comatose or time of recovery from mTBI) were recorded. Mice in the sham group (*n* = 6) were placed on the CHIMERA stage with anesthesia and without mTBI for the same 2–3 min duration. After gaining consciousness, mice were returned to their respective home cages. Based on preliminary experiments, we have found that mice with repeated mild injury tend to develop allodynia. Hence, the above procedure was repeated once a day for 4 days. All animal procedures were conducted in accordance with the NIH and AVMA guidelines and approved by the Uniformed Services University of the Health Sciences Institutional Animal Care and Use Committee (IACUC).

**Fig 1.**
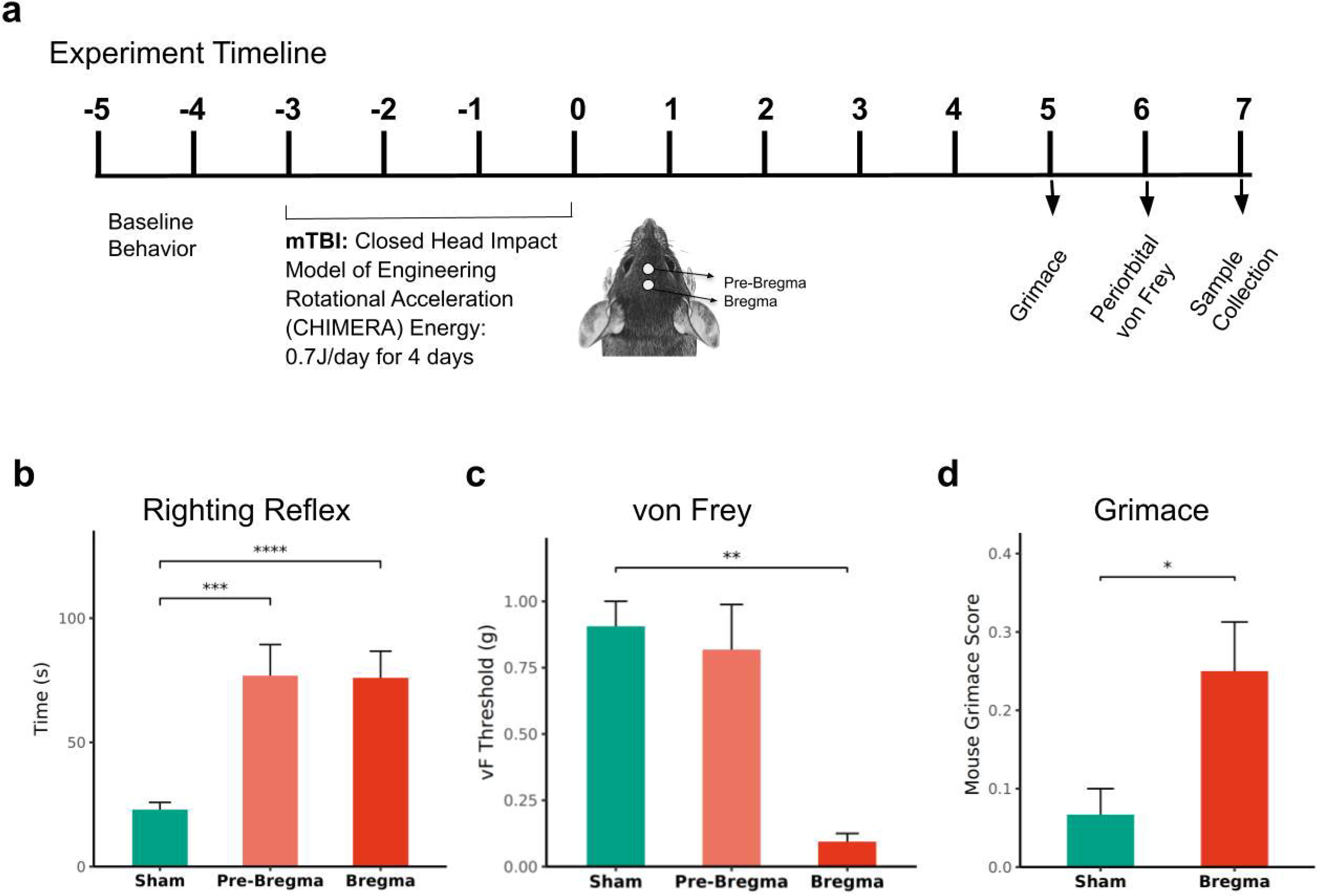
Behavioral assessment following mTBI in mice. a) schematic representation of the experimental procedure, with an illustration depicting the position of mTBI hits on the dorsum of the head and a timeline depicting behavioral measurement and sample collection. **b** Increased righting reflex, a measure of comatose state, in mice following mTBI in the bregma or pre-bregma positions. **c** Depending on localization of injury, mTBI induced differential peri-orbital sensitivity or mechanical threshold measured by von Frey filaments in the periorbital region. **d** Increased grimace score in mTBI mice observed 5 days after the last mTBI hit on bregma position. *, *p* < 0.05, **, *p* < 0.01, ***, *p* < 0.001 and ****, *p* < 0.0001. (*n* = 5–6/group, Mean ± SEM)

### Behavioral assessment

On the 6th day after the last TBI impact, mechanical allodynia was measured in the periorbital region using von Frey method. Mechanical threshold measurements were obtained using a simplified Chaplan up-down method [124, 125]. Prior to von Frey test, each mouse was tail restrained in a red tube and habituated for 10 min once a day for two days. On the test day, a series of calibrated von Frey filaments (Bioseb, France) ranging from 0.02 g to 2 g were applied to the midline periorbital region between the eyes. Each filament was perpendicularly placed on the midline region and pressed firmly with sufficient force to bend the filament. A positive response was defined as a rapid withdrawal of the head, or shaking of the head, or retrieving back or face wipe. A minimum of 20–30 s elapsed between filament tests.

To measure spontaneous pain response following mTBI, a separate cohort of mice was tested for grimace using mouse grimace score [40]. Prior to mTBI, mice were habituated in plastic animal enclosures (approximately 10 × 20 × 15 cm) for 15 min and on test days each mouse was tested for 10 min. Baseline and post-injury grimace scores were obtained from several still pictures and if needed, grimace was confirmed from video recordings. Of the 5 standard mouse grimace features, grimace such as periorbital tightening and altered ear positions were prominent in this mouse cohort and were used in the grimace analyses. In a separate cohort to determine long term effect of mTBI, grimace was measured on day 6, day 16 and day 22 post mTBI. All tests were performed between 10 AM and 3 PM. The experimenters were blind to the treatment conditions.

### Quantitative RT-PCR (qPCR)

To determine regional differences in gene expression following mTBI in pre-bregma or bregma positions (*n* = 5–6/group), fresh frozen samples from several brain regions were collected by cryosectioning at 250 µm using dissection mentioned above or using Palkovits punch method. We collected different pain-related brain regions, including Sp5C, PAG (described above), parabrachial nucleus/locus coeruleus (PB/LC), and thalamus. Because the parabrachial nucleus (PB) and locus coeruleus (LC) are in close proximity, both PB and LC were collected together during the Palkovits punch method. Further, to identify temporal variations in the gene expression profile, in a separate cohort (*n* = 6–7/group), fresh TG samples were obtained day1 and day 8 after the last mTBI. Total RNA was extracted using Zymo Research spin column as per manufacturer’s instructions. Because TG samples are highly myelinated, excess TRI Reagent was used to yield a good amount of RNA. RNA (1-2 µg) was reverse transcribed using Maxima First Strand cDNA Synthesis Kit, and qPCR was performed using primer sets for genes listed in in Table 1.

**Table 1.**
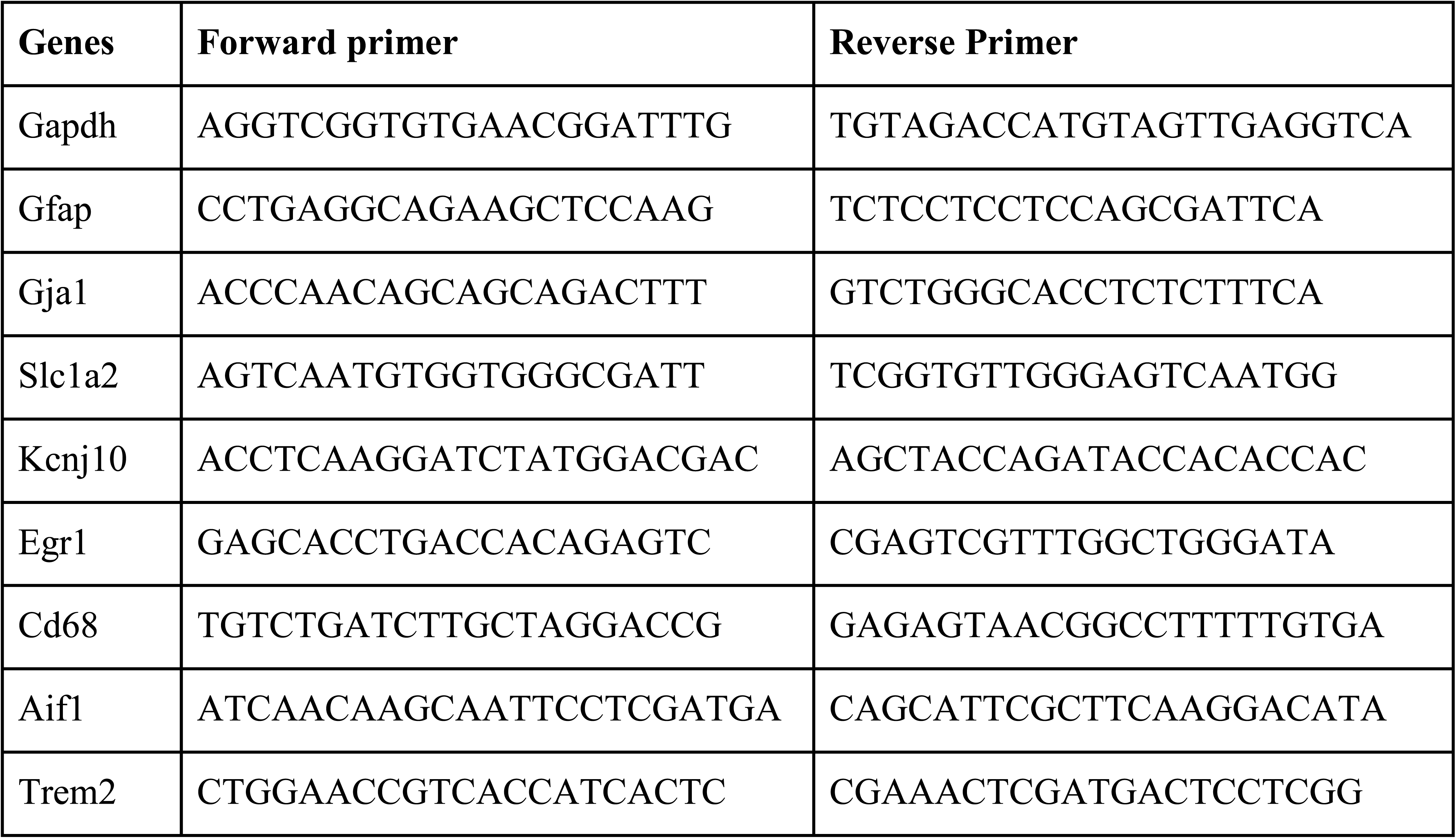
List of primes used in quantitative PCR.

### Bulk RNAseq and gene set enrichment analysis

Seven days following the repetitive mTBI, flash frozen TG (n=3/group), cryo-dissected Sp5C (n=3/group; bregma from -8.15 mm to -7.64 mm and medio-lateral 1.25 mm) and PAG (n=3/group; bregma from -3.4 mm to -4.96 mm and medio-lateral 0.65 mm) were processed for sequencing. RNA-seq was performed at GENEWIZ (Chelmsford, MA). RNA was extracted using Rneasy Plus Universal mini kit and RNA from TG, Sp5C and PAG samples had a RIN>8.4 (TapeStation) and concentrations were determined by Qubit RNA Assay. The following configuration was used for sequencing: Illumina HiSeq 2×150bp paired end sequencing on an average of 35M reads per sample. Deseq2 analysis was performed to identify differentially expressed genes using apeglm algorithm in lfcShrink function [43].

Gene set enrichment analyses (GSEA) and KEGG pathway analyses were performed using ‘clusterprofiler’ and curated mouse gene sets from ‘msigdbr’ libraries obtained from Molecular Signatures Database (MSigDB) were used to identify enriched gene sets in the mTBI group. Gene ontology of biology processing from C5 collections were used for GSEA. Dotplots for the GSEA were obtained using tidyverse’s ‘ggplot’ or ‘pathview’ and ‘enrichplot’ bioconductor packages.

Due to recent advances in single cell transcriptomics, there is an increased understanding of the TG cell specific gene expression in headache pathology and TG specific cell atlas [35,36]. Custom gene sets were built containing gene sets specific for TG cell populations [36]. These custom-built reference gene sets were then used for enrichment analyses to identify specific TG cell types involved in mTBI. We then also performed a meta-analysis to identify commonly expressed genes between the current mTBI dataset and different headache or injury models (induced by cortical spreading depression, inflammatory soup, spared nerve injury, facial lesion and scratch). Because these transcriptomic datasets are from single nuclear RNAseq, pseudobulk differential expression analysis was performed (as described in Harvard Chan Bioinformatics Core for scRNA-seq pseudobulk DE analysis) and Deseq2 tool was used as described above. For meta-analysis we used p-value combined approach using Fishers methods available in metaRNAseq or MetaVolcanoR packages.

### In situ hybridization and immunohistochemistry

Based on gene set enrichment analysis and differential expression data, selective genes were used for *in situ* hybridization using RNAscope Multiplex Fluorescent v2 assays (Advanced Cell Diagnostics, Inc, Newark, CA). The hybridization assay was performed according to the manufacturer’s instructions. Fresh frozen protocol was used for TG (12µm thick sections) and the fixed-frozen protocol was used for the Sp5C (15 µm thick sections) with protease treatment for 15 min and 30 min, respectively. Localization of markers of satellite glial cells such as Slc1a2 (Cat No. 441341), Gfap (Cat No. 313211) and Kcnj10 (Cat No. 458831) were used to identify the specificity of cell types in the TG. To identify the expression levels of mRNA (Gfap - Cat No. 313211) as well as protein in the Sp5C, immunohistochemistry was performed following in situ hybridization using manufacturer’s instructions. Localization of the GFAP was identified using Anti-GFAP (1:500 dilution; Dako Polyclonal Rabbit Z033429-2), followed by rinsing and incubation with a secondary antibody (Alexa-Fluor 488).

To determine inflammatory response by assessing microglial activation in the Sp5C, we used CD68 (1:300 dilution; eBioscience FA-11; rat anti-CD68) and Iba1 (1:500 dilution; Novus NB100-1028; goat anti-Iba1) in a separate assay. For assessing inflammation induced pain signaling (see below), we also performed calretinin immunostaining (1:500 dilution; Swant 6B3; goat anti-calretinin) in the Sp5C and co-immunostained with neurofilament L (1:5000, Encor; RPCA-NF-L-Degen) to define the boundaries of the superficial lamina.

Imaging of the tissue sections used for quantitative analysis was performed using Zeiss Axioscan Z1 Slide Scanner. Confocal images were obtained using the Zeiss 980 Confocal microscope. To identify the fine cell process, z-stack images at 1-2 µm intervals were acquired and maximal projection was applied to all images. Calretinin intensity measurement was performed using Profile View in Zen Lite 3.9.

### Statistical analysis

All analyses were performed using R 4.2.3 (https://www.R-project.org/). Packages in tidyverse library [42] were used for data processing and analyses. Behavioral and quantitative gene expression data among sham, bregma and pre-bregma mTBI groups were analyzed using mixed effect models or analysis of variance (ANOVA). Further, due to variability in between sections, fluorescence intensity within tissue sections was normalized. Quantitative cell counts or intensity of immunostains between sham and mTBI group were analyzed using stats package lm regression model or welsh test using lme4 package. Effects of mTBI were considered significant at an alpha level of 0.05. All datasets were tested for normality assumptions and when assumptions were violated, non-parametric test (Wilcoxon test) was used. All data were represented as mean ± standard error.

## Results

### Periorbital allodynia and spontaneous pain behavior in mice with repetitive mTBI

Acceleration head injury model by CHIMERA resulted in an increased righting reflex on each of the 4 mTBI days, which reflected an increased unconscious state. On average, irrespective of the hit position, the righting reflex of mTBI mice was ∼53 seconds more than the sham group (F(2, 93) = 7.7, p = 0.00079; Fig. 1b). On each of the four repetitive mTBI days, no significant difference was observed between bregma and pre-bregma hit groups (Additional File 1).

In contrast to the increased righting reflex, 6 days after the last mTBI, only mice that received repeated mTBI hits in the bregma position showed significantly reduced mechanical threshold in the periorbital region, compared to the sham group (F(2, 23) = 9.41, p = 0.001; Fig. 1c). Mice that received repeated mTBI in the pre-bregma position had an estimated coefficient of -0.08 g (p = 0.59), whereas the mTBI group in the bregma position had an estimated coefficient of -0.81 g compared to the sham group (p <0.01). Since pre-bregma position hit did not induce periorbital sensitivity, grimace, gene expression and histology were performed in mTBI mice that exhibited periorbital sensitivity from hit in the bregma position. In a separate cohort, where mice received repetitive hits in the bregma position, grimace was measured at 5 days post-injury. Spontaneous pain measured using grimace was significantly increased in the mTBI group compared to shams (F(1, 15) = 7.118, p = 0.01755; Fig. 1d). Further, a significant increase in grimace was noticed on week 2 and week 3 following mTBI (Additional File 2).

### Gene expression in the trigeminal pain regions following mTBI

Next, we used quantitative PCR to determine mRNA expression in several pain associated regions following mTBI in the pre-bregma or bregma positions. Recent evidence shows that Egr1, early growth response 1 - an immediate early gene, is associated with injury to ganglionic cells [35, 45]. To identify whether mTBI induced injury in the TG the expression of Egr1 was examined. Similar to mTBI mice (bregma hit) exhibiting mechanical allodynia, only the mTBI bregma group showed increased Egr1 mRNA expression in the TG (p = 0.018, Fig. 2a) compared to the sham group. Since gliosis and neuroinflammation are the primary sequelae to injury, we also examined generic astrocytes (Gfap) and activated microglia/macrophage (Aif1, Cd68, Trem2) markers in different brain regions including the Sp5C, PAG, PB/LC and thalamus. As anticipated, different regions had differential expression of these genes (Fig. 2b) and the expression levels of these markers were considerably higher in the bregma hit group (p = 1.48e-07) than the pre-bregma hit group (p = 0.00579), which were both greater than the expression in the sham group. In particular, the markers for microglia/macrophages were upregulated in the Sp5C compared to PAG, PB/LC and thalamus, while the astrocyte marker gene expression was highly upregulated in the PAG compared to Sp5C, PB/LC and thalamus.

**Fig 2.**
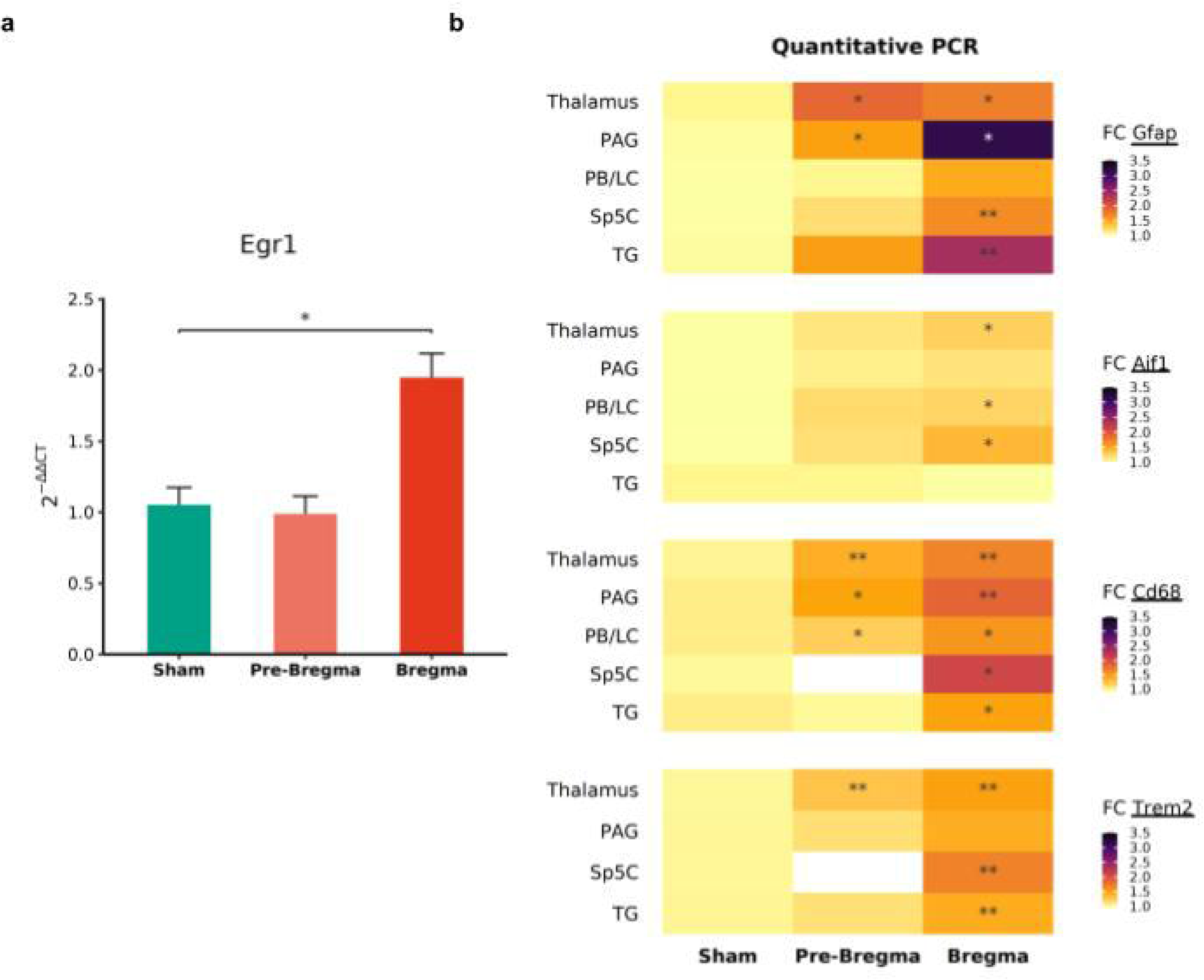
Quantitative gene expression in the TG and brain regions following mTBI on pre-bregma or bregma positions. a) Fold change (qPCR) in Egr1 (early growth response 1) mRNA expression in the TG 7 days post. Data (*n* = 6) in the bar graph are represented as mean ± standard error. *, *p* < 0.05. b) Heatmap representing qPCR fold changes (FC) in the gene expression of Gfap, Aif1, Cd68 and Trem2 in the Sp5C, PAG, PB/LC and the thalamus. *, *p* < 0.05, **, *p* < 0.01 (*n* = 5–6/group)

### Regional transcriptomics and gene set enrichment analysis following mTBI

Following the initial gene expression analyses, three pain related regions (TG, Sp5C and PAG) from bregma hit position were sequenced for subsequent analyses. As expected, transcriptomics data obtained from the TG, Sp5C and PAG revealed distinct gene expression profiles (Fig. 3a). More than 70% of the variance was observed between peripheral (TG) and central (Sp5C/PAG) regions. Whereas, within the brain, 21% variance was observed between the Sp5C and PAG. Similarly, the number of differentially expressed genes varied between regions (with log fold change > ±1, p-adjusted < 0.05 and base mean count >10). Specifically, differential expression of 4862 genes between TG and Sp5C, 5268 genes between TG and PAG and, within the brain, 2056 genes between Sp5C and PAG were noted.

**Fig 3.**
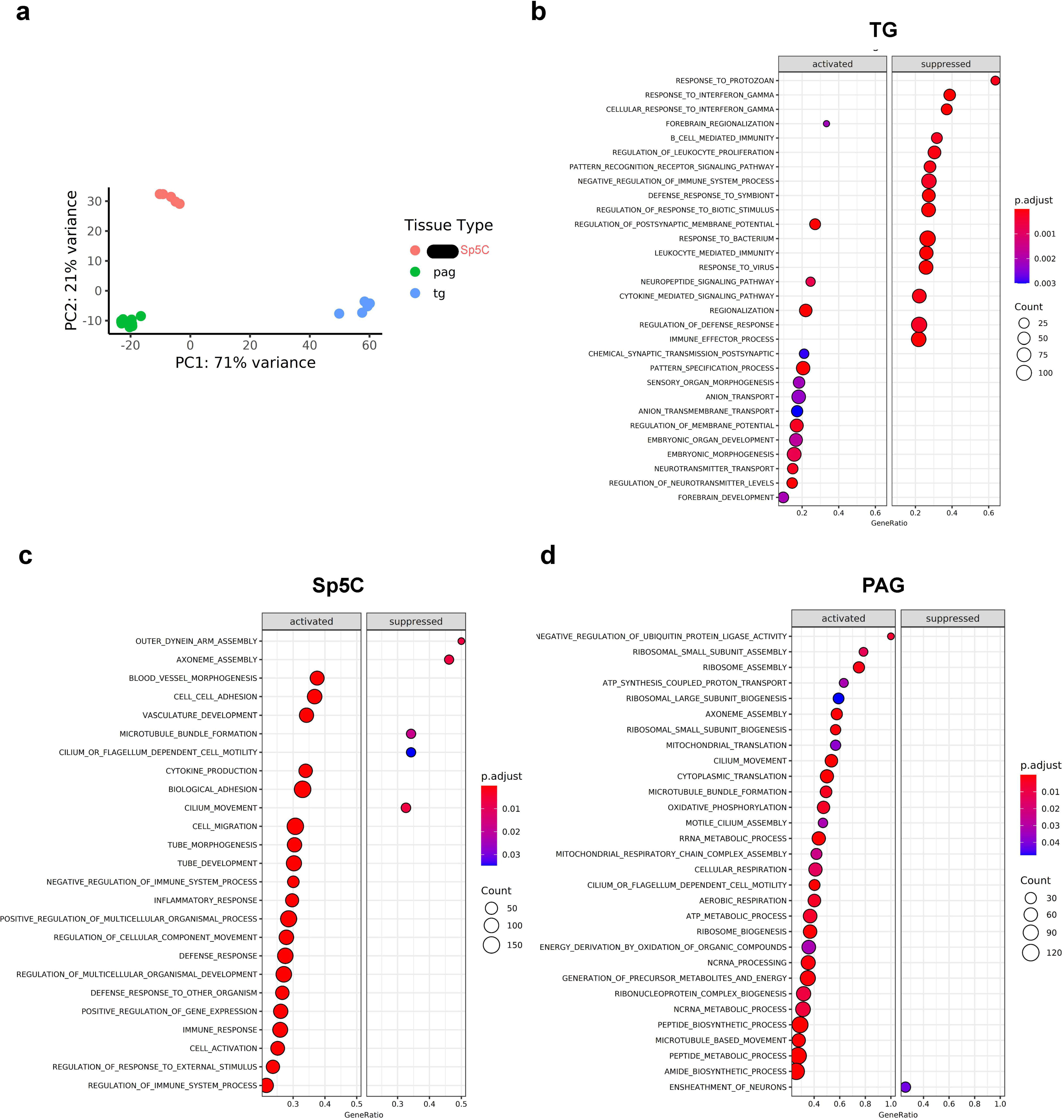
Transcriptome analyses from TG, Sp5C and PAG following mTBI. a) Principal component analysis of mouse transcriptome from TG, Sp5C, and PAG shows three distinct gene clusters belonging to the peripheral ganglion (TG), the medulla (Sp5C) and midbrain (PAG) regions. b) Dot plots of Gene Set Enrichment Analysis (GSEA) in the TG following repetitive mTBI, representing the 15 most differentially enriched biological process gene sets obtained from the C5 collection in the mouse molecular signatures database. Dot plots of GSEA analysis in the Sp5C (c) and PAG (d) of mTBI mice, representing the most differentially enriched biological process gene sets obtained from the C5 collection in the mouse molecular signatures database. Data used for PCA and GSEA were obtained from n=3/group/region.

Gene set enrichment analyses showed a distinct profile between TG and Sp5C or PAG following mTBI. Although gene sets involved in cell differentiation, tissue repair and remodeling were commonly expressed following mTBI, gene sets related to immune functions were suppressed in the TG but not in the Sp5C or PAG (Fig. 3b-d, Additional File 3). Particularly, following mTBI, gene sets from biological processes of gene ontology showed decreased enrichment (negative sign of normalized enrichment score) of the following GO terms: B cells, T cells, leukocytes, cellular defense response, response to virus, defense response to bacteria, and response to interferon gamma in the TG. Nonetheless, postsynaptic regulation, regulation of ion transport, neuropeptide signaling pathway, and neural precursor of cell proliferation were enriched in the TG of mTBI mice.

In contrast to the TG, with an exception of a few, the majority of the gene sets in the Sp5C of medulla and the PAG of the midbrain, were enriched 7 days following mTBI. The only gene sets with negative enrichment score in the Sp5C following TBI were related to microtubule organization which includes axoneme assembly, cilium movement, flagellum dependent cell motility, and microtubule bundle formation. In the PAG, only the gene set related to ensheathment of neurons was negatively enriched. Full list of enriched gene sets obtained from GSEA analyses from all the regions were presented in Additional File 3.

### mTBI induced TG specific gene set enrichment analysis and meta-analysis

A GSEA analysis using custom gene sets specific for TG cell types [35,36] showed that non-neuronal cells were highly enriched compared to neuronal cells following mTBI (Fig. 4a). Particularly, a positive enrichment score was identified for fibroblast, myelinated schwann cells (p < 0.05) and satellite glial cells (p = 0.18). While immune cells and neurons (somatostatin, neurofilament cluster - NF 1-3, non-peptidergic, cLTMRs) specific gene sets showed negative enrichment (p < 0.05). Enrichment of fibroblast and satellite cells was consistent with other headache models, such as migraine induced by cortical spreading depression (CSD) or inflammatory soup (IS) [36].

**Fig. 4.**
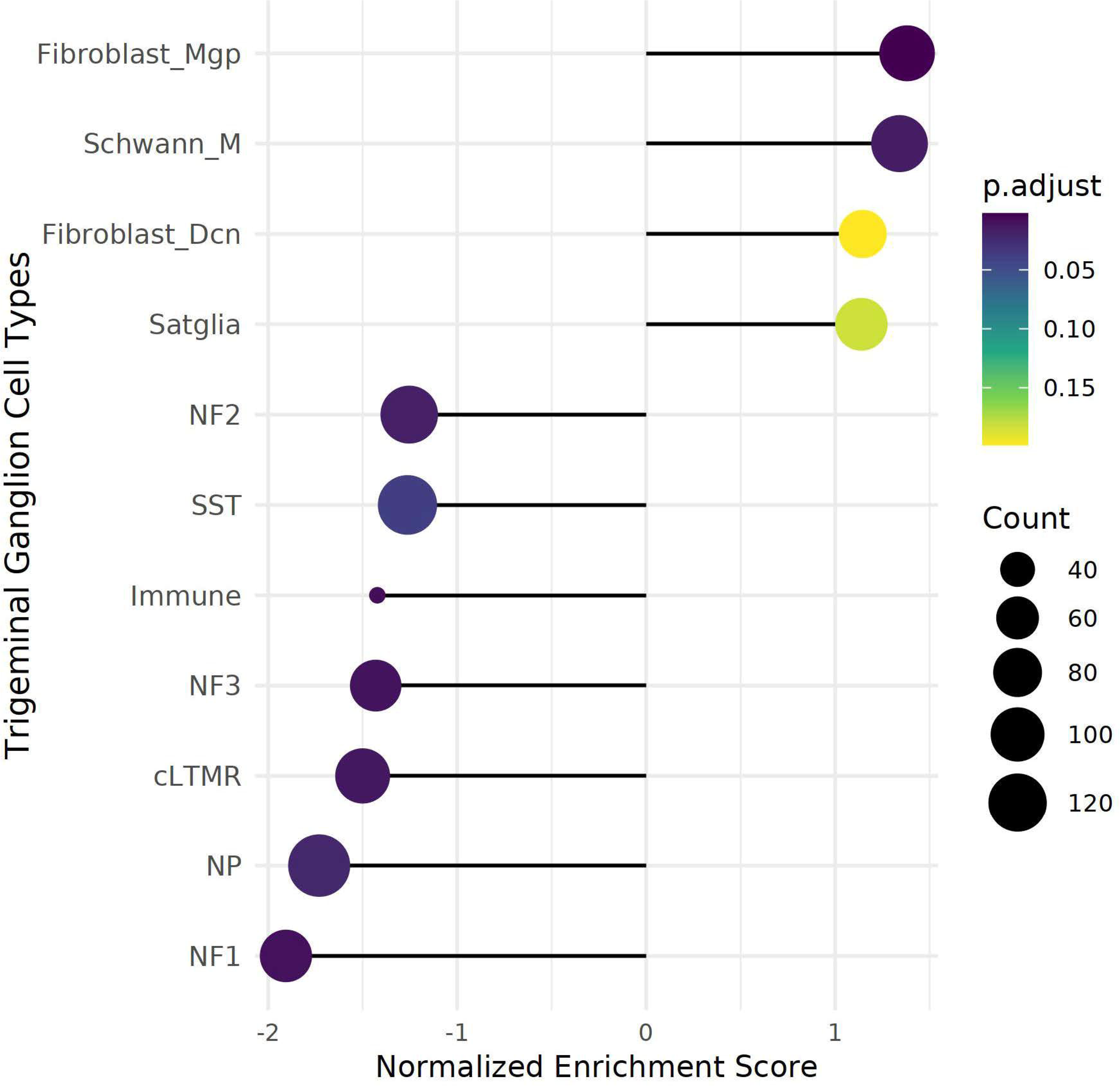
Comparison of the TG transcriptome of mTBI mice with TG cell atlas. a) Dot plot representing enriched gene clusters of different cell types in the TG. Non-neuronal gene clusters have positively normalized enrichment scores (p-adjusted values <0.2) and neuronal gene clusters have negative normalized enrichment scores (p-adjusted <0.05). Abbreviations: cLTMRs - C-fiber low threshold mechanoreceptor (LTMR); fibroblast_Dcn - DCN+ meningeal fibroblasts; fibroblast_Mgp - MGP+ meningeal fibroblast; NF1, neurofilament+ A-LTMR enriched for A-beta-field; NF2, neurofilament+ A-LTMR enriched for A-beta-RA and A-beta-field; NF3, neurofilament+ LTMR enriched for A-delta; NP - non-peptidergic nociceptor; Schwann_M, myelinating Schwann cells; SST, somatostatin positive pruriceptors.

Secondly, the overlapping genes between mTBI and other headache or TG injury models were identified using meta-analysis. Because TG samples were collected 7 days following mTBI, only minimal fold changes were noted in the current transcriptomic dataset. Thus, to identify overlapping genes between mTBI and different TG pathology models, a less constraint fold change was used in the current dataset. Since the sequencing datasets from spared nerve injury (SNI), facial lesion and scratch injury [35] were based on TG neuronal population, the number of overlapping genes were reasonably lower compared to those obtained from all cell population in the CSD and IS models [36]. Nonetheless, a majority of the overlapping genes were upregulated (Table 2). A few genes such as Lmo7 (LIM domain 7), involved in protein-protein interaction, overlapped among all models including mTBI, and Nts (neurotensin) overlapped among mTBI, CSD, spared nerve injury, facial injury and scratch. Kccn3 (Potassium Calcium-Activated Channel Subfamily N Member 3) overlapped between mTBI, CSD and IS and belongs to TRPM8 (Transient receptor potential melastatin subtype 8) neuronal cluster. Neuronal injury marker, Atf3 - ATF/CREB family of transcription factors, was overlapped between mTBI, SNI, facial lesion and scratch injury, reflecting Egr1 expression in mTBI. Non-neuronal cell markers particularly for schwann cells, satellite glia, fibroblast and vascular endothelial cells were commonly expressed between mTBI, CSD and IS. Interestingly, satellite glial markers such as Gfap, Iqgap2 (cytoskeleton interacting protein), Slc6a6 (sodium and chloride-ion dependent transporter) were shared among mTBI, CSD and IS. Only a few shared genes (Hmx1, Itga11 and Fhit) were down regulated among mTBI, CSD and IS and were related to distinct neuronal clusters (Table 2).

**Table 2.**
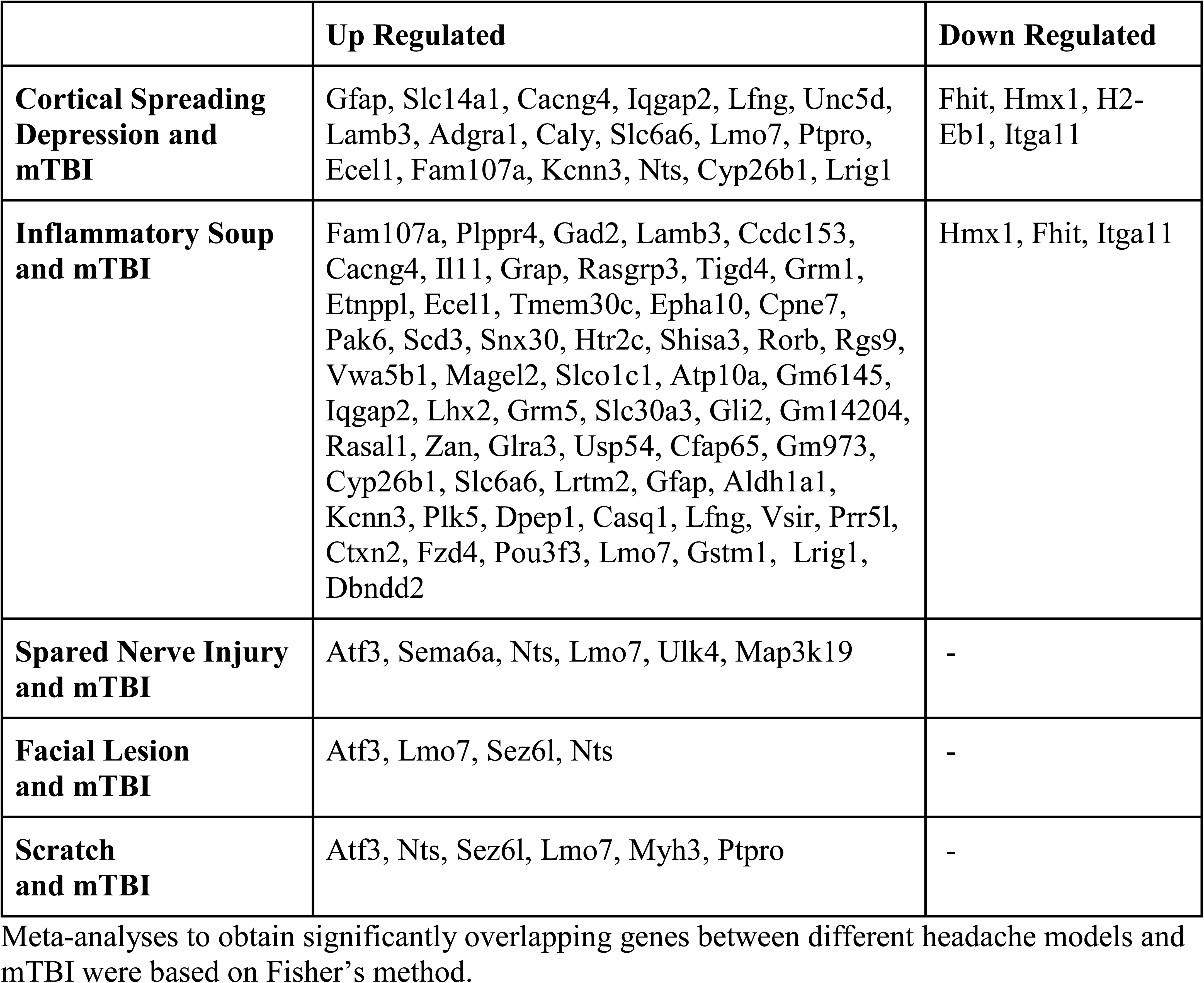
Comparison of overlapping genes between mTBI and other TG pathology models.

Although spinal dorsal horn and Sp5C share some similarities in the rexed laminae appearance, due to unique somatotopy and circuitry, Sp5C have quite distinct neurophysiological properties [44]. Since, Sp5C specific transcriptome is not available yet, we utilized the brainstem astrocyte specific gene sets [45] and found that astrocyte specific gene ontology gene sets were enriched in the mTBI (Additional File 4).

### Validation of gene expression in the TG

Transcriptomics data identified that gene sets associated with ion channel transport and satellite glial cells were positively enriched in the mTBI mice. Moreover, satellite glia cells have gained attention in the past several years due to their involvement in the pain mechanisms [20,21]. Therefore, we focused on the specific glia markers, Gfap (glial fibrillary acidic protein), Kcnj10 (potassium inwardly rectifying channel subfamily J member 10), Gja1 (gap junction protein, connexin 43), and Slc1a2 (solute carrier family 1 - glial high affinity glutamate transporter). In a separate cohort (n = 7-8/group), quantitative gene expression of satellite glia markers revealed that Gfap and Slc1a2 (Fig. 5a) were upregulated one day after the last mTBI and continued to show increased expression 1-week post mTBI (p<0.01). Gja1 was slightly increased on day 1 (p = 0.0525), but was significantly upregulated at 1-week post-injury (p = 0.0339, Fig. 5a). Using multiplex in situ hybridization, mTBI mice showed increased expression of these markers in satellite glia surrounding neurons of the TG (Fig. 5b).

**Fig. 5.**
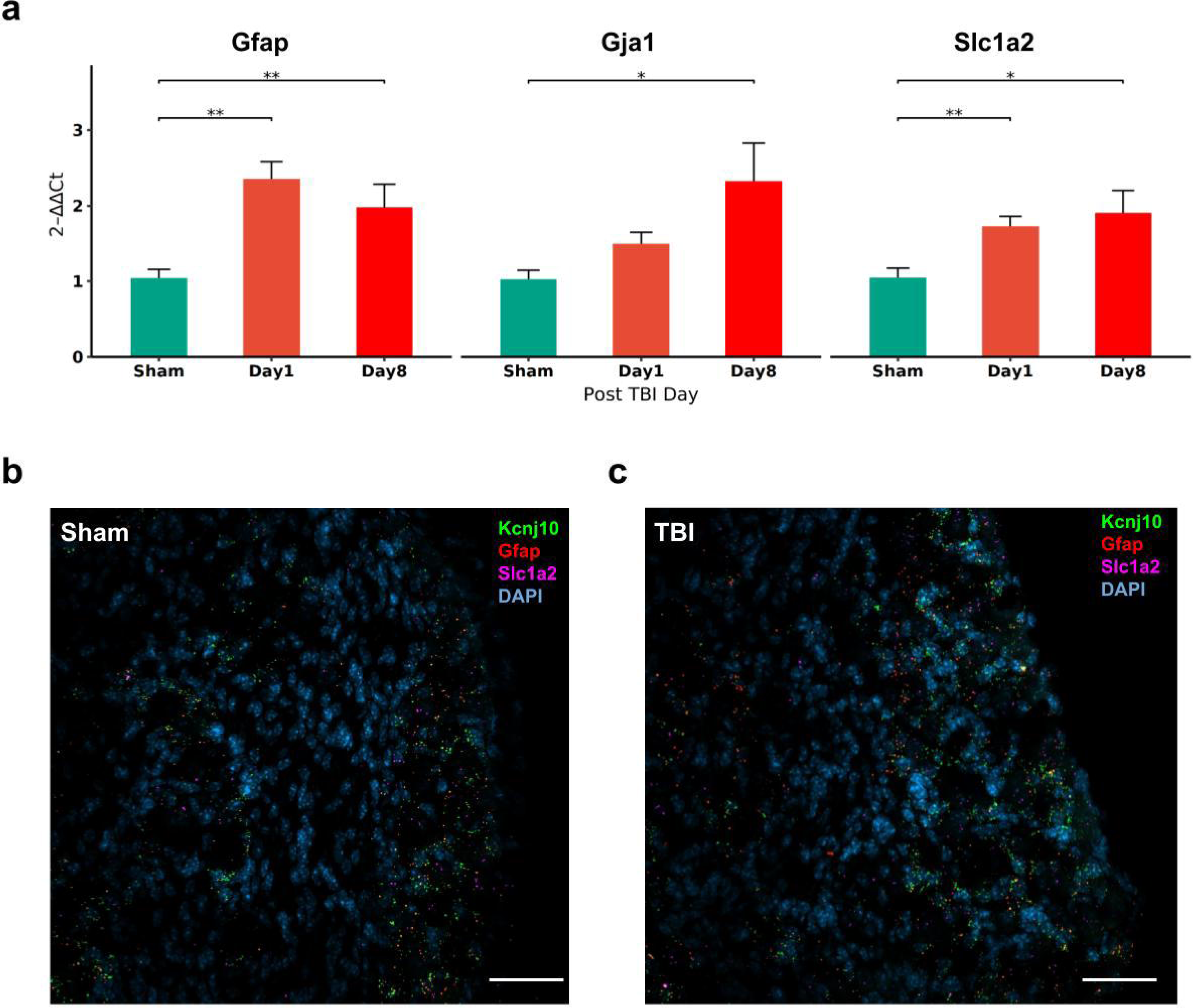
Mild TBI induced gene expression of satellite glial markers in the TG. a) Fold change (qPCR) in gene expression of satellite glial markers (Gfap, Gja1 and Slc1a2) in the TG of mTBI mice. Bar graphs show mRNA expression on day 1 and day 8 after the last mTBI. Data (n=5-6/group) in the bar graph are represented as mean±standard error. *, p<0.05, **, p<0.01. b) Localization of mRNA expression of Kcnj10 (green), Gfap (red) and Slc1a2 (magenta) in the TG of sham and mTBI mice obtained at 7 days post injury. Scale bar = 50µm.

### Neuroinflammation in the trigeminal nucleus caudalis

Trigeminal nucleus caudalis showed a wide range of gene expression changes in the transcriptomic data. Importantly, GO terms such as inflammatory response, immune response, cytokine production and regulation of immune system processes were highly enriched in the Sp5C. Diffuse nature of mTBI injury and innervation of primary trigeminal afferents, to the Sp5C, between trigeminal subnuclei and other brainstem regions [46–48], have prompted our investigation of activated microglia using immunohistochemistry in different regions of rostral-caudal extent of the trigeminal Sp5C (Fig. 6). Based on the area and cell number, a significant proliferation of CD68+Iba1+ cells in the white matter, particularly in the spinocerebellar tract (sc, p = 3.01e-11; Fig. 6a-c) was noted, while a moderate increase in the tectospinal tract (ts, p = 0.0038) and pyramidal tract (py, p = 0.0013) was observed. In the gray matter, the gracile nucleus (Gr) was found to have significantly increased expression of CD68+Iba1+ cells (Gr, p < 0.001). Inter reticular nucleus showed a moderate increase in microglia (IRt, p = 0.0407) and no differences were noted in the reticular nucleus (MdV, p=0.623). Although no significant difference was found in the rostral Sp5C (p = 0.192) and caudal Sp5C (p=0.4), CD68 intensity within the Iba1+ cells in Sp5C was significantly higher in the mTBI group (Fig. 6d, P < 0.0001). Interestingly, these Iba1+ cells were in close association with GFAP+ cells/fibers (Fig. 7f and also see below). Furthermore, morphological differences in microglia were noted in the white matter tracts, gracile nucleus and Sp5C (Additional File 5).

**Fig. 6.**
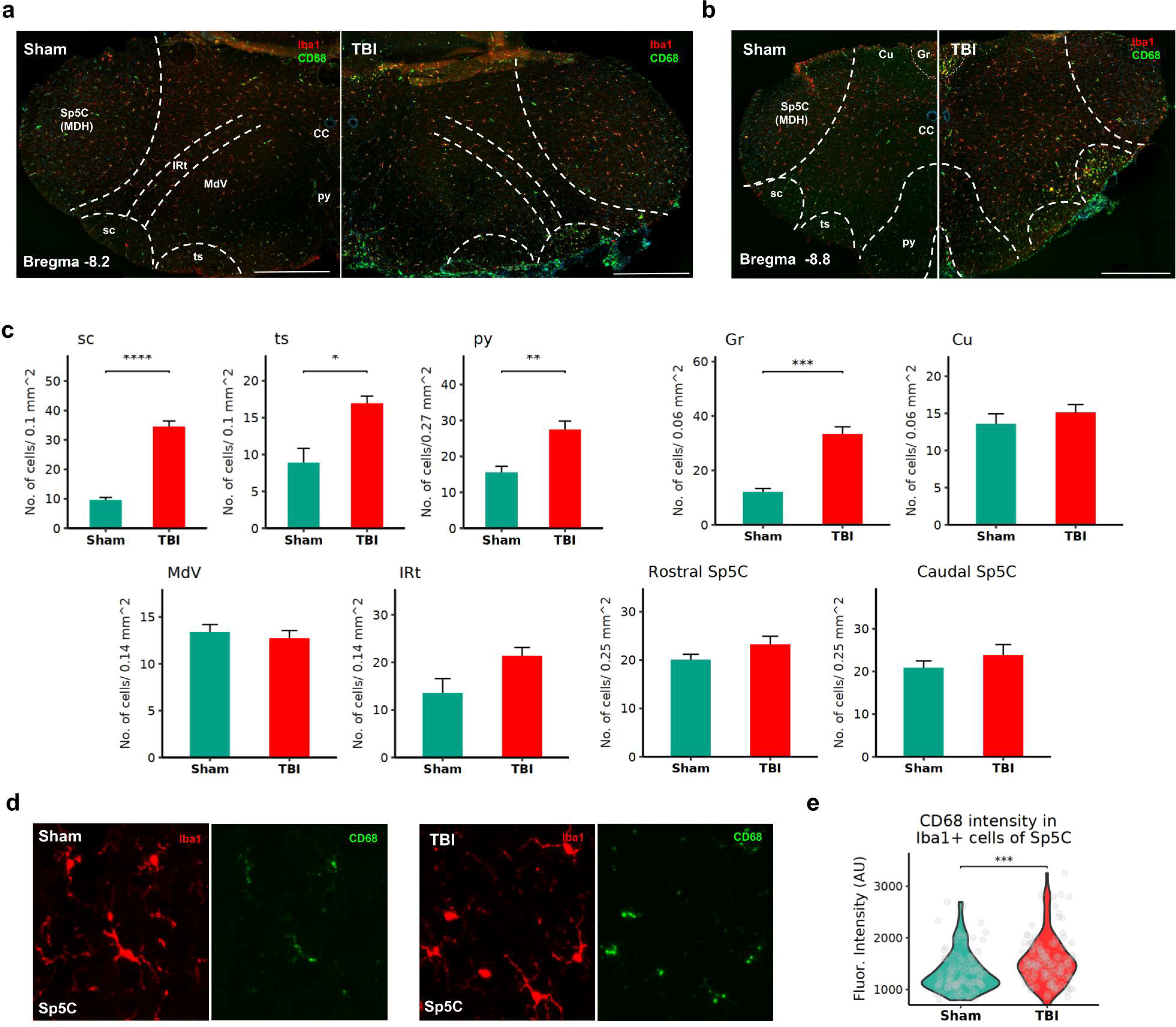
Differential expression of inflammatory markers using immunohistochemistry in the trigeminocervical complex of mTBI mice. a) Representative images from sham and mTBI mice showing immunostaining for CD68 in, Iba1 + cells in the caudal Sp5C (Bregma coordinate ∼ -8.6) and (b) the rostral (Bregma coordinate ∼ -8.1). c Increased CD68/Iba1 + cell count, most notable in the fiber tracts and in the gracile nucleus of mTBI mice. d and e Increased CD68 intensity within Iba1 + cells in the Sp5C of mTBI mice compared to the sham group. *n* = 10–15 cells/section; *n* = 3 mice /group. Abbreviations: Cu— cuneate nucleus; Gr—gracile nucleus; IRt—Intermediate reticular nucleus; Mdv—Medullary reticular nucleus, ventral part; py—pyramidal tract; sc—spinocerebellar tract; Sp5c (MDH)— spinal trigeminal nucleus caudalis (medullary dorsal horn); ts—tectospinal tract. *, *p* < 0.05, **, *p* < 0.01, ***, *p* < 0.001 and ****, *p* < 0.0001. (*n* = 5–6/group, Mean ± SEM)

**Fig. 7.**
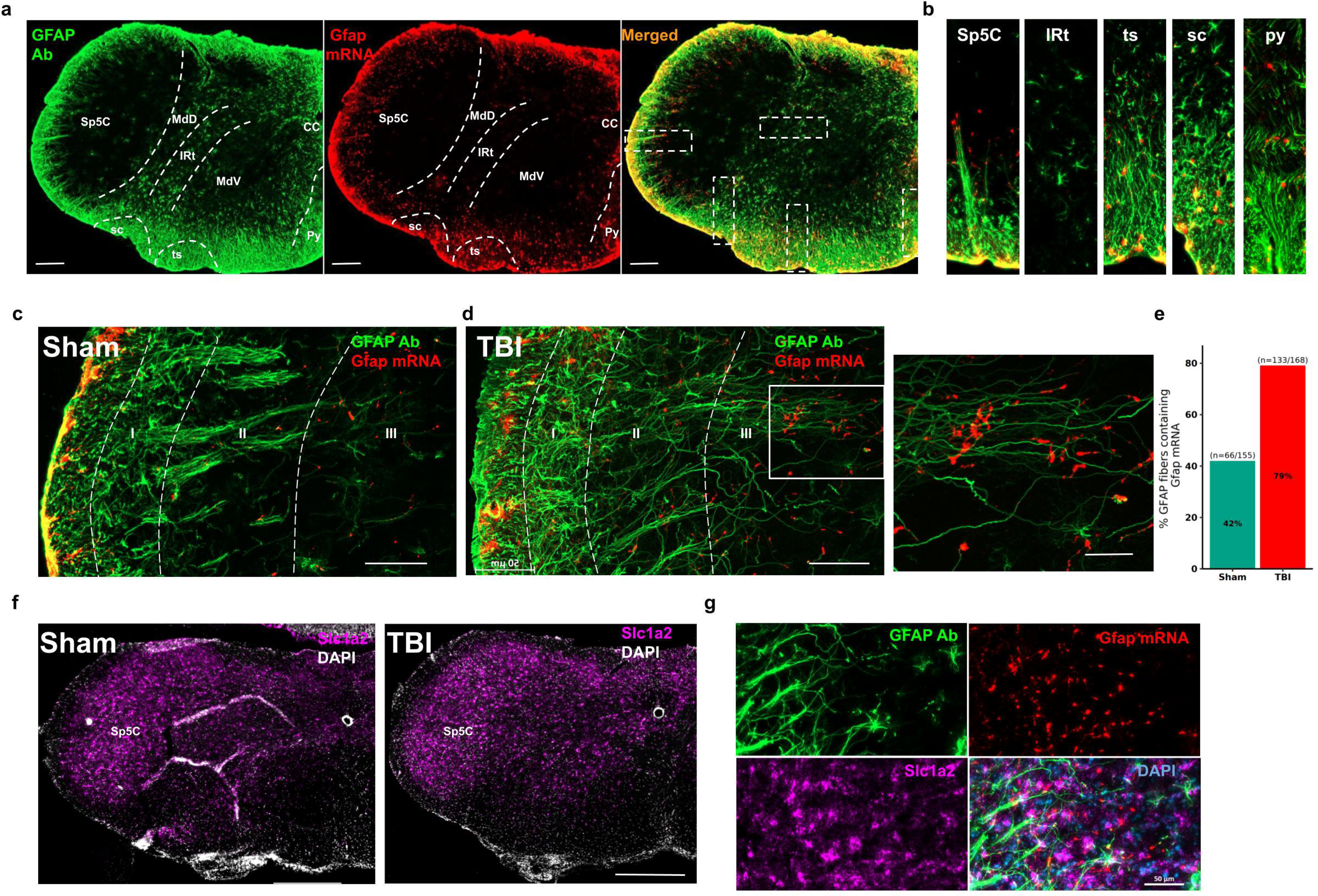
Reactive astrogliosis in brainstem of mTBI mice 7 days following injury. Reactive astrogliosis in brainstem of mTBI mice 7 days following injury. a) An example brainstem section at the Bregma level -8.2 stained with GFAP (green, immunohistochemistry) and Gfap (red, in situ hybridization). Scale bar = 200 µm. b) Insets in Fig. 7a representing differences in Gfap mRNA/GFAP antibody (Ab) staining in various brainstem regions. Representative confocal images from a sham (c) and a mTBI (d) mouse showing colocalization of Gfap mRNA (red, in situ hybridization) on GFAP + astrocytes (green, immunohistochemistry) in the Sp5c. Scale bar = 50 µm. Inset of 7d from a mTBI mouse Sp5C shows increased Gfap mRNA expression in the processes of GFAP + astrocytes in lamina III of Sp5C. Scale bar = 20 µm. e) Graph representing percent Gfap mRNA expression in GFAP + fibrous astrocytes in the Sp5c of sham (*n* = 155 fibers) and mTBI (*n* = 168 fibers) groups. f) Close proximity of GFAP-ir fibers or cells to Iba1/CD68 + cells in different regions of the brainstem are shown in white arrows. GFAP-ir fibers or cells that are not in close contact with Iba1/CD68 + cells are shown in purple arrows. g) Representative images showing localized expression of Slc1a2 in the Sp5C of sham and mTBI mice. Scale bar = 500 µm. h) Presence of GFAP + astrocytes (green, immunohistochemistry) in close proximity to Slc1a2 mRNA puncta (purple, in situ hybridization) in the lamina III of Sp5C are shown in white arrows. Scale bar = 50 µm. IRt— Intermediate reticular nucleus; Mdv—Medullary reticular nucleus, ventral part; py—pyramidal tract; sc—spinocerebellar tract; Sp5c (MDH)-spinal trigeminal nucleus caudalis (medullary dorsal horn); ts—tectospinal tract

Because astrocytes specific gene sets were also enriched in the Sp5C transcriptomic data (Additional File 4) and known to be involved in inflammation [49,50], in parallel to the TG, we also investigated glia expression in the Sp5C of mTBI mice. Similar to gene expression changes in the TG, Gfap mRNA level in the Sp5C was upregulated 1-week post-injury (Fig. 2b and Fig. 7c-e). In contrast to the TG, Gfap mRNA expression was found colocalized on long astrocytic processes identified by immunohistochemistry (Fig. 7a-c). Particularly, a fold increase in Gfap mRNA expression was observed at the long processes of GFAP+ astrocytes in mTBI mice (Fig. 7e). Although Gfap mRNA was present in the superficial layer of Sp5C and increased in mTBI (Fig. 7c and 7d), due to high density of astrocyte soma in the superficial layer we were unable to quantify Gfap mRNA in GFAP+ soma. Immunostaining also showed that astrocytes in the Sp5C have a distinct expression pattern as they transverse through deep layers of the Sp5C. Compared to the reticular formation (Fig. 7b), only sparse presence of protoplasmic astrocytes were noted in the Sp5C (Fig. 7a-b and Additional File 6). Moreover, in some sections, Gfap+ astrocytes processes were bundled in the sham group while these astrocytes processes were separated in the mTBI group. Although GFAP+ processes traversed from lamina 1 and deeper into lamina 3 (Fig. 7d insert), no significant differences were found in the length of GFAP+ processes in the Sp5C between sham and mTBI groups.

Due to the increased interest in the astrocyte-microglia interaction in neuroinflammation [51], we also performed triple immunohistochemistry targeting GFAP, Iba1 and CD68+ cells in the brainstem sections. Irrespective of the morphology of GFAP (i.e. either fibrous or protoplasmic astrocytes), a substantial proportion of the GFAP+ fibers or cells were in close contact with Iba1/CD68+ cells (Fig. 7f). Interestingly, lamina 1 of the Sp5C had more GFAP+ fibers in contact with Iba1/CD68+ cells (Fig. 7f - inset 1) than lamina 2 or 3 (Fig. 7f - inset 2-3). Other regions in the brainstem, such as reticular formation and white matter tracts also expressed GFAP+ cells and fibers, respectively, in close proximity to the Iba1/CD68+ cells (Fig. 7f - inset 4-6).

### Cellular mechanisms of pain modulation in the trigeminal Sp5C

Since we found increased reactive astrogliosis in the Sp5C of mTBI mice and glutamatergic signaling is well known to be involved in the pain modulation, we next investigated the changes of the glutamate transporter (Slc1a2 gene product of GLT1/EAAT2) in the trigeminal region, Sp5C. Interestingly, the Slc1a2 expression was well defined in the gray matter, particularly in the Sp5C. In contrast to our expectation, the expression of Slc1a2 was ubiquitous without clear boundaries and no identifiable difference was observed between sham and mTBI mice (Fig. 7g). In order to test whether GFAP+ fibers in the Sp5C were in close proximity to Slc1a2 expression, we performed in situ hybridization of Gfap and Slc1a2 combined with GFAP immunohistochemistry, and found that the majority of GFAP+ fibers containing Gfap mRNA were in close proximity to Slc1a2 mRNA expression in lamina 2 and 3 of Sp5C in mTBI mice (Fig. 7h).

Further to identify the causal relationship between cellular/molecular changes observed above and headache-like symptoms, we sought to characterize calretinin, a marker involved in allodynia associated with inflammatory pain [52]. We found a reorganization of calretinin immunoreactivity in the Sp5C (Fig. 8a, 8b) where the peak intensity of the calretinin immunoreactivity shifted from outer lamina II to lamina I in the Sp5C of mTBI mice (p = 0.0136). Moreover, we also identified that the number of calretinin cells differed between rostral-caudal extent of Sp5C and was higher in the mTBI mice (Fig. 8c). Although, the number of calretinin cells were lower in the rostral compared to caudal Sp5C, a significant increase was noted in the rostral Sp5C (p= 0.004) and a non-significant increase was noted in the caudal Sp5C (p = 0.1935).

**Fig. 8.**
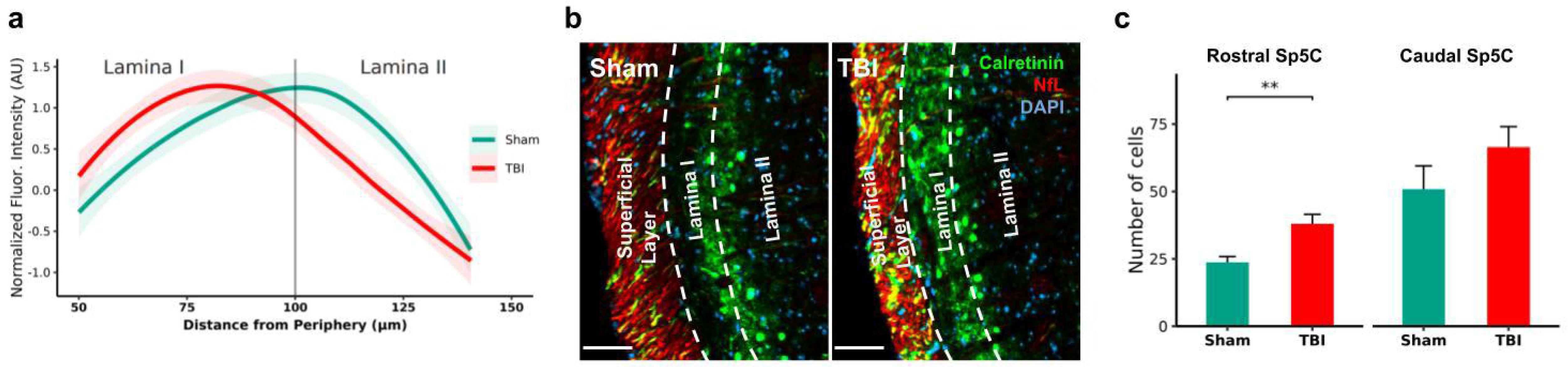
Calretinin in the Sp5C of mTBI mice potentially involved in trigeminal pain. a) Reorganization of calretinin-ir was observed from outer lamina II to lamina I of mTBI mice (n=3; Bregma coordinate ∼8.2). Full extent of this reorganization is shown in Additional File 7. b) Representative images from sham and mTBI mice at Sp5C show the shift in calretinin-ir from lamina II to lamina I. Scale bar = 50 µm. c) Increased number of calretinin-ir neurons in the rostral Sp5c compared to the caudal Sp5c of mTBI mice. **, p<0.01.

## Discussion

We found that non-invasive mTBI using the acceleration method induced comatose state as well as headache-like symptoms, which resemble the hallmark comorbid conditions of TBI observed in clinical populations [8,53–56]. Distinct expression profiles and increased inflammation in the pain associated regions following mTBI showed that there was a concerted effort of different brain regions involved in multimodal components of pain [57,58], thereby experiencing both mechanical allodynia as well as spontaneous nociceptive pain response.

### Repetitive mTBI induced allodynia and grimace

In the clinical population, mild head injury has higher prevalence for developing PTH [6,11,59]. Similar to clinical cases of mTBI in both nonhuman primates and humans [8,54,56], where patients typically lose consciousness for less than 30 minutes, our mouse model exhibited unconsciousness for a few minutes post-injury. It has been reported that the number of injuries correlates with an increase in persistent headache symptoms [6] and brain pathology [60]. This set the precedent for repetitive head injury in our model where mice developed periorbital sensitivity and grimace over time. In line with this observation in mice, decreased mechanical threshold is one of the characteristics of patients experiencing post-traumatic headache [61,62]. Particularly, here we have shown that there are differences in the site of injury on head where mTBI mice exhibit distinct mechanical pain thresholds and varying degrees of inflammatory gene expression in the brain. Although mice were in comatose state after receiving mTBI either in the bregma or pre-bregma position on the dorsum of the head, only mice that received mTBI in the bregma position displayed periorbital sensitivity and grimace thereby mimicking symptoms similar to post-traumatic headache. This could potentially be related to injury to the trigeminal nerve terminals surrounding the head [18,63–65] and these results might explain the heterogeneity involved in mTBI induced symptoms. Effects of mTBI on grimace several days after injury pointed to the fact that mTBI induced by the acceleration method had clinical manifestation where headache symptoms were observed after injury [66]. The profound innervation of the head by the TG, as well as its first and second order connections to the Sp5C, PB, and PAG, suggests a potential role for these regions in the chronification of PTH [57,67,68].

### Repetitive mTBI induced differences in gene expression

Having the proof of concept that periorbital sensitivity and grimace become prominent after repetitive mTBI, we investigated gene expression changes in the TG and several other pain related regions. Specifically, sensory information from the trigeminal system is relayed to the ascending pain regions, including PB/LC [65,69,70], PAG [71–73], and the thalamus [74]. Similar to existing literature on the difference in the location of impact and regional differences in brain pathology [75,76], we found that non-invasive traumatic injury to the head, only at the bregma region, exhibited prominent changes in Egr1 gene expression in the TG and elevated gene expression of astrocytic and inflammatory markers in the Sp5C, PB/LC, PAG and the thalamus. This enhanced pathology of mTBI in the bregma region was likely due to the proximity of injury to both coronal and sagittal sutures of the cranial vault, neurovasculature, and the biomechanics involved in the trauma [65,77,78]. Together, in our mTBI mouse model, injury to the bregma position of the head has increased TG/brain pathology that likely led to increased headache-like symptoms characterized by periorbital sensitivity and grimace.

To comprehensively assess gene expression following mTBI, we performed deep sequencing of the TG, Sp5C, and PAG in mTBI (bregma position) and sham groups. Due to the mild injury model used in this study and sequencing performed in samples collected 7 days post mTBI, fewer than 10 genes resulted in log2 fold change of greater than 1 in the tissues examined. Nonetheless, gene sets enrichment analyses provided an opportunity to understand co-regulated genes from these samples. At 7 days following repetitive mTBI, we found the peripheral TG had a distinct gene expression profile compared to the central Sp5C and PAG. The immunosuppressive state of the TG following mTBI might render it vulnerable to subsequent infection or injury and possibly leading to hyperexcitability of sensory neurons under certain pathogenic conditions [79–82] or even by migraine-like triggers provided several days post injury [83–86]. In contrast to TG, the majority of gene expression in the Sp5C and PAG were upregulated following mTBI. The only difference between the Sp5C and PAG was gene sets involved in the microtubule formation and cilium movement, where these genes were downregulated in the Sp5C and upregulated in the PAG. This difference could be due to the proximity of brainstem Sp5C compared to the PAG, where severe axonal stretching may have led to microtubule disassembly [87,88].

To further understand the cellular mechanisms of mTBI induced injury in the TG, we took advantage of recently published TG cell atlas [36]. Using the gene enrichment analyses, it was identified that non-neuronal gene clusters were highly enriched compared to neuronal gene clusters. Enrichment of fibroblast and satellite glial cells was consistent with other mouse headache models (inflammatory soup or cortical spreading depression), but the enrichment of vascular cells shown in the inflammatory soup model [36] was not observed in our mTBI model. We also found that gene sets related to myelinated schwann cells were highly enriched in the mTBI and this could be of importance as schwann cell proliferation has been shown in other nerve injury models [89,90] and might be involved in trigeminal pathology. Further, meta-analysis also revealed a distinct set of genes from this study significantly overlapped with severe injury models such as infra-orbital transection (spared nerve injury), facial lesion or scratch, inflammatory soup or cortical spreading depression models of headache. Injury markers [36] such as Atf3, and Sema6a overlapped between mTBI and severe injury models (spared nerve injury, facial lesion and scratch) reflecting that mTBI induced injury to the TG. Interestingly, Nts, neurotensin, was overlapped in the majority of the injury or headache model, and it has been well documented that neurotensin modulate neuropathic pain [126]. A number of genes related to schwann cells, fibroblast, satellite glia and vascular endothelial cells significantly overlapped between mTBI and headache models (CSD and IS), indicating that non-neuronal cells of the TG have a major role in headache pathology. A few ion channel markers, such as Kccn3 (potassium Calcium-Activated Channel Subfamily N Member 3) and Slc6a6 (sodium and chloride-ion dependent transporter) overlapped between mTBI, CSD and IS, where Kcnn3 has been associated with migraine pathology [127]. Nonetheless, less than 50 genes overlapped between mTBI and other models. As such, mTBI induced a distinct gene expression profile compared to other TG pathology models. Collectively, increased susceptibility of TG and distinct gene expression profile in the TG following mTBI could explain the headache heterogeneity following mTBI in clinical populations [2] and the challenge in translatability using animal models [32]. Nevertheless, non-neuronal gene clusters, particularly, satellite glia have gained particular interest for their potential role in headache pathology [20,36].

### Glial activation in the trigeminal regions following mTBI

In light of recent evidence, satellite glial cells show a remarkable functional heterogeneity in sensory ganglion [20,91, 21,92]. Here we focused on the expression of specific satellite glial cell markers following mTBI. Consistent with the literature, increased gene expression of Gfap, Gja1 and Connexin 43 (Cx43) suggested that mTBI induced increase in satellite glial activity could function as a protective mechanism [93], or to facilitate pain [20,91] as the expression of these genes continued to increase through the post-injury days. In contrast to the existing evidence in other model systems [94–96], we found mTBI increased Kcnj10 expression in the satellite glia surrounding sensory neurons. This might suggest a protective role of glia in the TG through buffering extracellular potassium [97–100] where satellite glial cells could prevent further depolarization of sensory neurons. Additionally, the increased expression of glutamate transporter, Slc1a2, in the mTBI group signifies that glutamate signaling is involved in mTBI induced neuropathic pain as shown in other injury models [101,102]. Since, there is high diversity in satellite glial cells [20,91], increased expression of specific markers found in this study suggests that satellite glial cells of the TG may play a diverse role in mTBI whose precise function in pain is yet to be determined.

Compared to the satellite glia cells in the TG, glia in the Sp5C possess a distinct morphology and may have distinct functional roles [103–105]. Due to regional differences in the gene expression pattern [45] and unavailability of trigeminal Sp5C specific transcriptional cell atlas, we applied gene sets (e.g. astrocytes) specific for the brainstem and identified that astrocytes were enriched in the Sp5C. Astrocytes in different lamina of the spinal cord dorsal horn are known to have specific functions, including modulation of nociception and facilitation of tactile information [106]. Interestingly, the morphology of Sp5C astrocytes closely resembles the interlaminar astrocytes present only in the primate outer cortices [107,108], or radial glial cells and, recent evidence shows that brainstem/cortical astrocytes have proliferative/migratory potentials [45]. Increased reactive gliosis or hypertrophied nature of Gfap+ fibrous astrocytes in the mTBI and the presence of Gfap mRNA in the end processes of these fibrous astrocytes in the Sp5C suggest that these astrocytes are dynamically modulated in the brainstem long after injury. Similar results in the Sp5C were reported in partial ligation of the infraorbital trigeminal nerve [37], where elevated astrocytes persisted even months after injury [76]. Although morphology of GFAP+ astrocytes are different in the spinal dorsal horn compared to Sp5C [109], because of the close proximity to Slc1a2, functionally, GFAP+ fibers in the Sp5C could modulate neurons via glutamate signaling involved in neuropathic pain [45,110–113]. In line with this, astrogliosis in the Sp5C has been shown to be involved in the development [114] and maintenance of pain [115]. Although we have not tested long term characteristics of reactive gliosis, Sp5C astrocytes may play a major role in the development of headache-like symptoms following mTBI and could be a target of treatment in chronic pain [116].

### Potential involvement of inflammation in mTBI induced headache

Due to the extensive damage caused by the acceleration mTBI model to the brainstem, it was expected that injury related pathology might extend beyond Sp5C. Microglial proliferation was more prominent in white matter tracts (spinocerebellar/tectospinal/pyramidal tracts) than in gray matter areas. While the role of microglia in white matter tracts is suggested to involve phagocytosis of degenerating axons, as seen in spinal cord injury [117], their function in gray matter may differ. For instance, activated microglia in the gracile nucleus, associated with touch and proprioception, might contribute to pain symptoms, as seen in preclinical studies linking TBI to plantar tactile hypersensitivity [83]. Although we did not assess plantar tactile sensitivity in this study, the presence of activated microglia in the gracile nucleus indicated a potential role for inflammation in the paratrigeminal region.

Additionally, reactive astrogliosis in various brainstem regions, often in close proximity to microglia, suggests potential astrocyte-microglia interactions influencing neuroinflammation and pain modulation [51]. The increased expression of microglial markers in the Sp5C seven days post-injury, combined with the extended reach of astrocytic fibers, hints at a possible recruitment of astrocytes by microglia during neuroinflammation [118]. This interaction could potentially modulate glutamate signaling in the Sp5C, given the proximity of astrocytic fibers to Slc1a2. The widespread presence of Slc1a2 throughout the Sp5C [119], indicates ongoing glutamate signaling in healthy mice for facial proprioception, suggesting that increased reactive astrocytes in mTBI might further influence this signaling pathway. Consequently, the combined effects of reactive astrocytosis and elevated microglia activation could potentially regulate pain within the trigeminal system.

Since we observed increased inflammatory response in the Sp5C and its neighboring region in the brainstem, we sought to elucidate underlying neural mechanisms of pain. Calretinin-positive interneurons, a population known for their involvement in inflammation-induced pain [52], increased in number and reorganize within the superficial laminae of the Sp5C following mild TBI. This pattern closely resembles with those observed in the spinal dorsal horn after peripheral nerve injury [120]. Calretinin interneurons have been implicated in the activation of ascending pain signals via projections to the parabrachial nucleus leading to chronification of pain [121]. Considering the predominant excitatory nature of these interneurons and their role in pain propagation [122,123], it is possible that reorganization from lamina II to lamina I and the increase in the number of calretinin-ir neurons following mTBI may contribute to the enhanced allodynia detected one week post-injury. Moreover, such recruitment would also explain an increase in microglia and astrocytes in the lamina I of the Sp5C to further augment ascending somatosensory pain signals. Further investigation into the interaction between glia and calretinin interneurons is warranted.

## Conclusions

Overall, mTBI in mice developed periorbital sensitivity and grimace days after injury thereby exhibiting headache-like symptoms. While widespread inflammatory markers were observed in several pain-related regions, transcriptomic analyses revealed that head injury induced distinct pathogenic gene expression profiles in peripheral TG and central Sp5C, where glia have distinct functional roles. Thus, glia in the trigeminal regions appear to modulate headache-like symptoms and offer an avenue for further research on treatment targets for PTH following mTBI.

## Availability of Data and Materials

The underlying data and computational scripts employed in this study will be shared upon request. Mild TBI induced differentially expressed gene from each tissue region (TG, Sp5C and PAG) are available in Additional file 3. Data from Yang et al., 2022 for custom gene sets used in GSEA and meta-analysis can be found in GEO GSE197289 and the supplementary tables Table S4, Table S12 (https://ars.els-cdn.com/content/image/1-s2.0-S0896627322002288-mmc5.xlsx and https://ars.els-cdn.com/content/image/1-s2.0-S0896627322002288-mmc13.xlsx) of the manuscript and data from Nguyen et al., 2019 used for comparison of injury models can be found in GEO GSE131272 and the supplementary file 3 (https://doi.org/10.7554/eLife.49679.019) of the manuscript.

## Abbreviations

mTBI: Mild traumatic brain injury
Sp5C: Trigeminal nucleus caudalis
TG: Trigeminal ganglion
PAG: Periaqueductal gray
IRt: Intermediate reticular nucleus
Mdv: Medullary reticular nucleus, ventral part
py: pyramidal tract
sc: spinocerebellar tract
ts: tectospinal tract
Iba1: Ionized calcium binding adaptor molecule 1
CD68: Cluster of differentiation 68
GSEA: Gene set enrichment analysis
CHIMERA: closed-head impact model of engineered rotational acceleration
Gapdh: Glyceraldehyde 3-phosphate dehydrogenase
Gfap: Glial fibrillary acidic protein
Gja1: gap junction alpha-1
Egr1: Early growth response factor 1
Aif1: Allograft inflammatory factor 1
Kcnj10: Potassium inwardly rectifying channel subfamily J member 10
Slc1a2: solute carrier family 1 (glial high affinity glutamate transporter), member 2

## Ethics approval and consent to participate

All animal procedures were conducted in accordance with the Office for Laboratory Animal Welfare (OLAW), National Institutes of Health (NIH) and American Veterinary Medical Association (AVMA) Guidelines for the Euthanasia of Animals guidelines and approved by the Uniformed Services University of the Health Sciences (USUHS) Institutional Animal Care and Use Committee (IACUC).

## Consent for publication

Not applicable.

## Authors’ contributions

GN and YZ designed the experiments. GN conducted the experiments, analyzed the data and wrote the manuscript. GN and YZ interpreted the data and revised the manuscript. Y.Z. acquired the research funding. All authors have read and agreed to the submitted version of the manuscript.

## Acknowledgements

We would like to thank Jie Wen, MD, Ph.D., for performing the von Frey testing. We also thank Amanda Fu, MD, and Laura Tucker, MS at the USUHS Preclinical Behavior and Modeling Core for assistance with mouse procedures, Dennis McDaniel, Ph.D., and Fritz Lischka, Ph.D., at the USUHS Biomedical Instrumentation Core for Imaging, and Gemini for text editing. We gratefully acknowledge the Student Bioinformatics Initiative (SBI) at the USUHS for providing crucial high-performance computing resources. We also extend our thanks to Andrew Frank, MS, Clifton Dalgard, Ph.D., and Matthew Wilkerson, Ph.D., for their insightful discussions and feedback on our bioinformatics approaches.

## Disclaimer

The statements and data presented in this manuscript belong to the authors of the manuscript and do not reflect the views of the Department of Defense or the Uniformed Services University of the Health Sciences and the The Henry M. Jackson Foundation for the Advancement of Military Medicine, Inc.

## Additional Files

**Additional File1**. Righting reflex in each of four mTBI days (n=6/group).

**Additional File 2**. Bar graphs representing grimace at 6, 16- and 22-days post mTBI (dpi) in bregma position (n=4-5/group).

**Additional File 3.** Enriched Go terms and differentially expressed genes from three different regions (TG, Sp5C and PAG) following mTBI.

**Additional File 4**. Dot plot representing astrocyte specific enriched gene sets in the Sp5C following mTBI.

**Additional File 5**. Distinct expression pattern of CD68+, Iba1+ cells in the brainstem. Fiber tracts, specifically spinocerebellar tract (sc), show a distinct morphological profile compared to the microglia in the gray matter (Sp5C, Gr, Mdv). Gr - gracile nucleus; Mdv - Medullary reticular nucleus.

**Additional File 6**. Increased presence of GFAP+ fibrous astrocytes in the Sp5C compared to the paratrigeminal regions. Rare appearance of protoplasmic astrocytes in the Sp5C and prominent protoplasmic appears in the reticular formation in both Sham and mTBI. IRt - Intermediate reticular nucleus.

**Additional File 7.** Representative images from sham and mTBI mice showing calretinin-ir neurons in the lamina I of Sp5C. Dotted white lines represent the Ib4+ staining. Atlas plate at the level of ∼ -8.3 from bregma. Scale bar = 200 µm.

