## Supplementary figures and images for "Distinct expression profile reveals glia involvement in the trigeminal system attributing to post-traumatic headache"

### Additional File 1

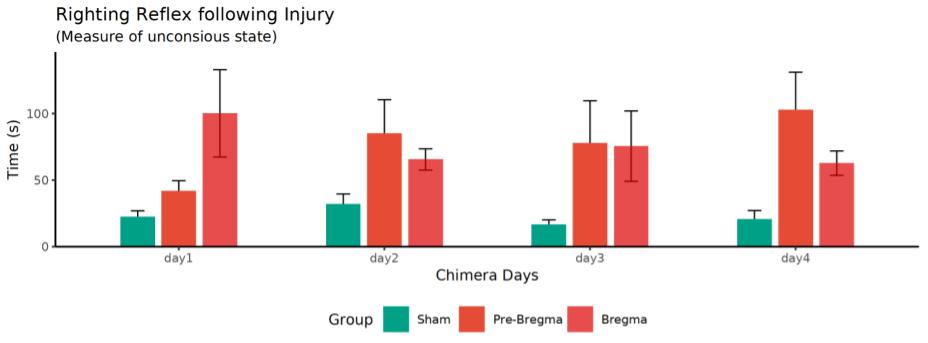

### Additional File 2

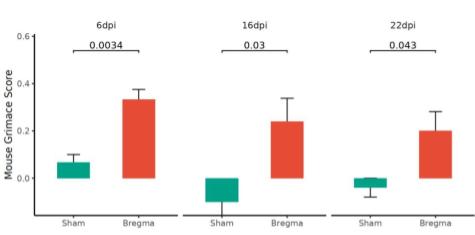

### Additional File 4

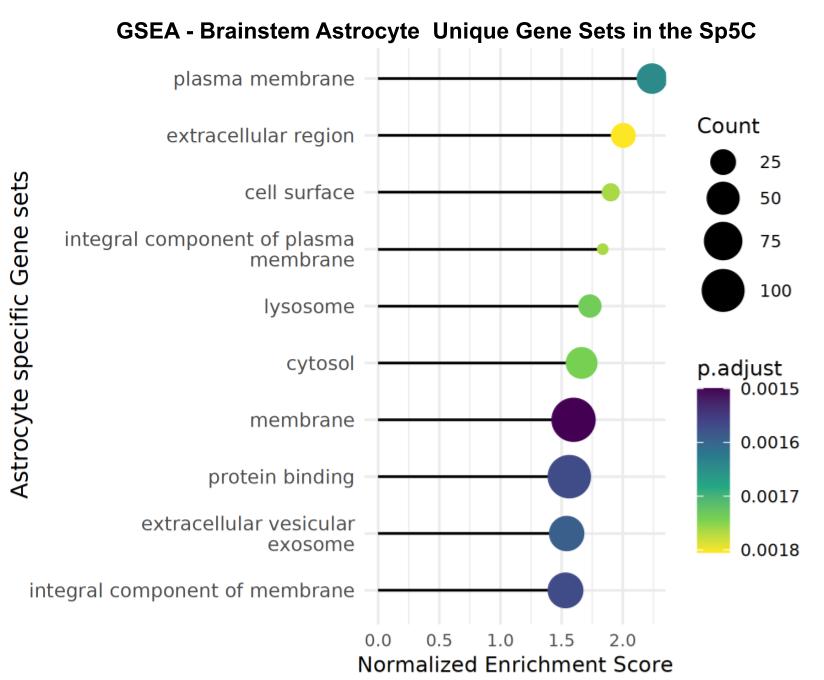

### Additional File 5

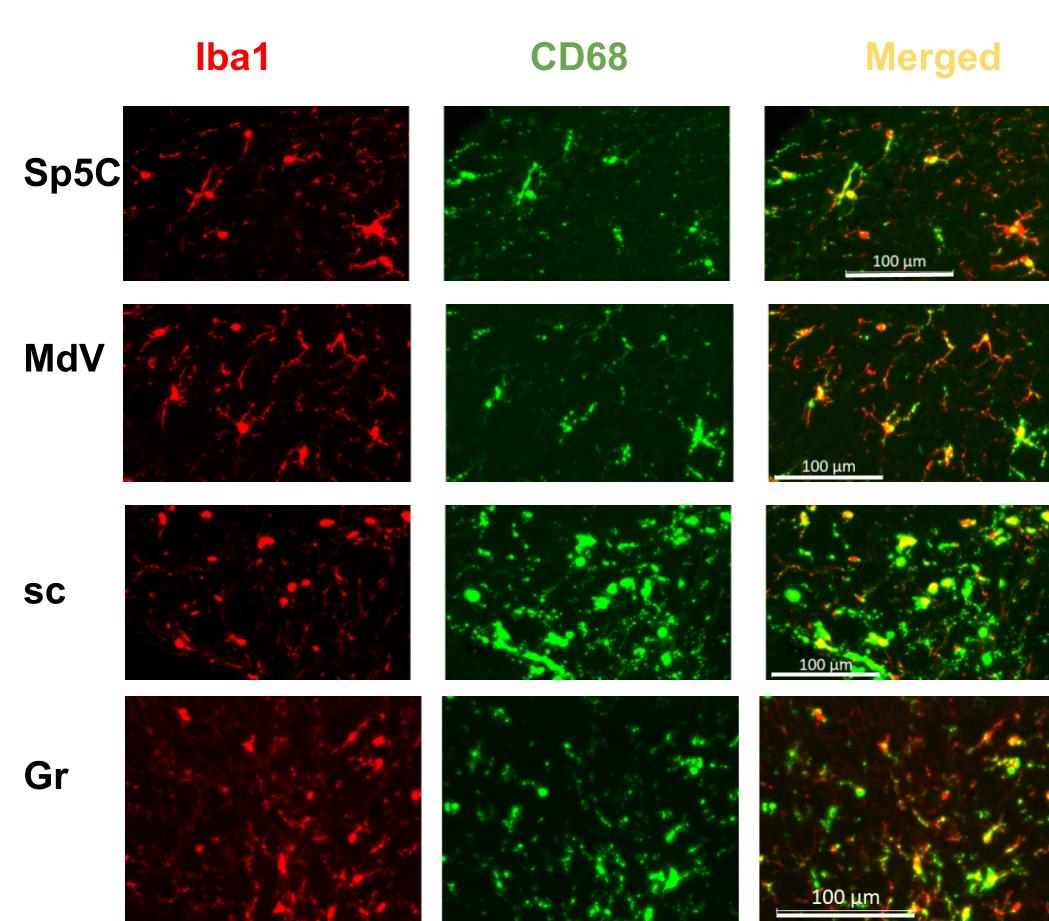

### Additional File 6

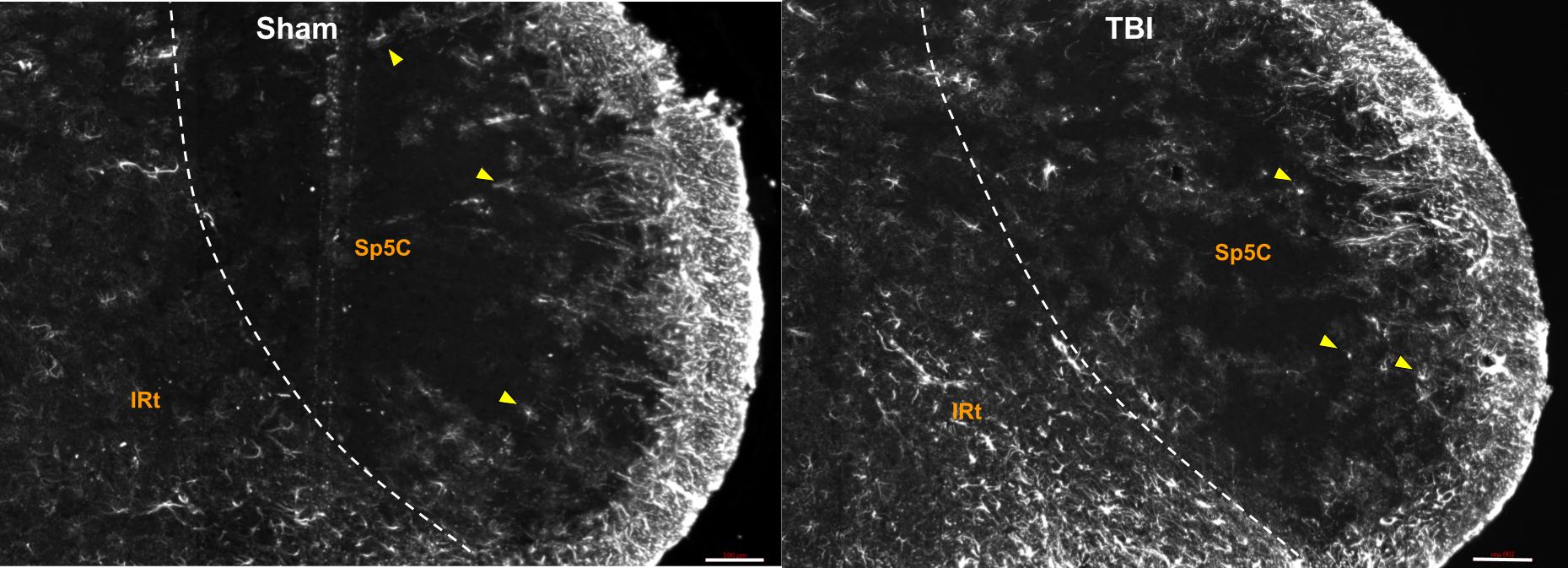

### Additional File 7

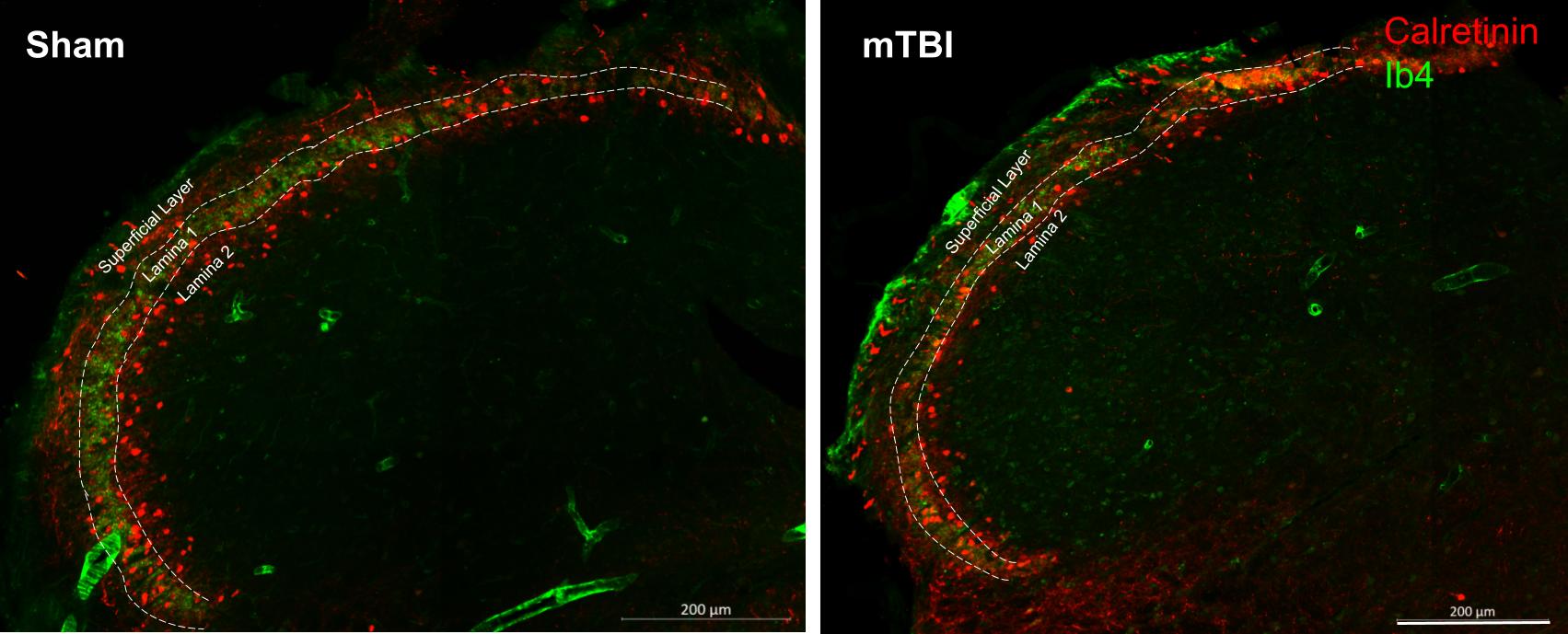
